# Why are spontaneous bio photons observed only in living organisms? The key role of the proton current in ultra-weak photon emission

**DOI:** 10.1101/2023.09.15.557955

**Authors:** Jerzy J. Langer, Marcin Langer

**Affiliations:** Laboratory for Materials Physicochemistry and Nanotechnology, Faculty of Chemistry, Adam Mickiewicz University in Poznań, Uniwersytetu Poznańskiego 8, Department of Physics of Functional Materials, Faculty of Physics, Adam Mickiewicz University in Poznań, Uniwersytetu Poznanskiego 2, 61-614 Poznań, Poland

## Abstract

Light emission from living things is still at the centre of scientific interest. Ultra-weak photon emission (UPE) in the range from 300 to 900 nm has been discovered in living cells and organisms, including the human body ^1^. In general, so-called bio photons are attributed to life.

Our recent studies on protonic p-n junction formation and light emission from electrically powered protonic p-n junction systems suggest, that UPE can be generated by excitations owing to proton current flow in living cells and sub-cellular structures (e.g. mitochondria), just like it is done in the case of laboratory protonic light-emitting diodes (H^+^LED) ^2,3^.

While the emission of higher energy bio photons (above 3 eV, 200-420 nm wavelength) is mainly caused by radicals and reactive oxygen species (ROS) ^1, 21, 32, 36, 37, 38^, lower energy bio photons (below 3 eV, at 420 -1000 nm wavelength) should be associated with the excitation of the protonic system as a result of the flow of the proton current (discussed in this paper). We expect this to have important biomedical implications for diagnosis and therapy using UPE ^36, 37^. The similarity of H^+^LED and UPE spectra (Fig. 2) allows the use of protonic H^+^LED as a new broadband light source, ideally suited to mitochondria-oriented low-intensity light therapy ^37^.

Our results explain why spontaneous biophotons (UPE) are observed only in living organisms, tissues and cells ^21^. This is due to the constant flow of protons in the active ATP synthase/ATPase ^23^ and in the mitochondria in general ^25-28^, which is necessary both for life and for the emission of light (observed as bio photons).

## Introduction

Mysterious emission of light from living things - ultra-weak photon emission UPE (also called bio photons) - is still at the forefront of scientific interest ^1^. Our recent studies suggest that UPE can be generated by excitation of the protonic system in living cells and sub-cellular structures (e.g. mitochondria), just like in the laboratory protonic light-emitting diode (H^+^LED) ^2, 3^.

In 1978 Peter Mitchell was awarded the Nobel Prize for the biologically oriented proton-based chemiosmotic theory. Since then, many studies have been conducted to explain what drives this mechanism ^7-13, 16-19, 22-24^. There are still many unanswered questions, including new aspects and phenomena ^26^.

Our experiments on protonic p-n junction formation and light emission from electrically powered protonic p-n junction systems, provide a basis for further research in the field combining light emission and proton flow. Just in our laboratory, a unique, first light emission (electro luminescence) induced in aqueous systems with a protonic p-n junction was observed, Fig.1. We also defined processes during light emission based on proton flow and excitations of protonic systems ^2, 3^.

**Fig. 1.**
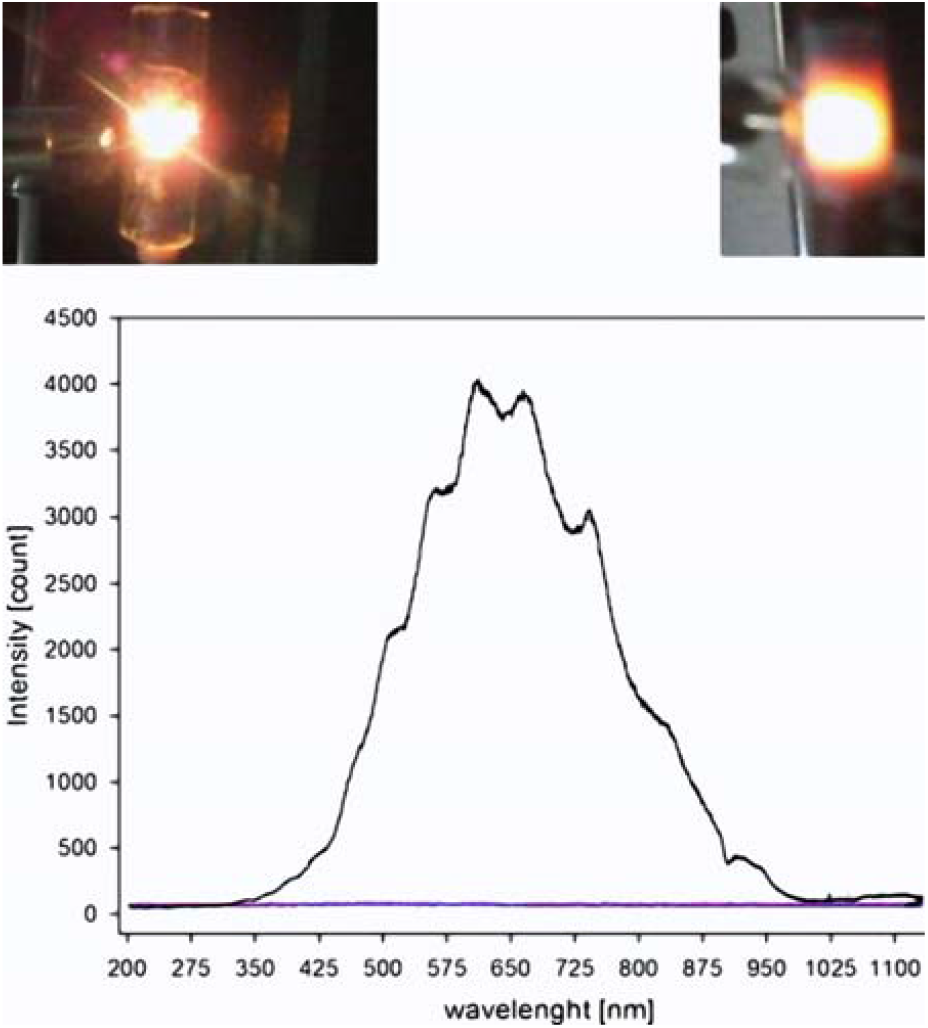
Emission spectrum of a protonic LED (H^+^LED) at 16V and 0.8 A. Insert: H^+^LED in action, protected by a glass tube (left) and by Teflon tube (right) ^2, 3^.

## Results and discussion

This is a new concept in light generation - protonic LEDs (H^+^LED) - based on protons instead of electrons. New light emitting devices H^+^LED are of potential importance for photonics, optoelectronics, photochemistry, and now we think - also for biology.

In particular, our research leads to the conclusion that bio photons (ultra-weak photon emission, UPE) can be generated by excitation of the protonic system in living cells at the cellular and sub cellular level due to the flow of proton current in the protonic p-n junction, similarly to the protonic diode H^+^LED (compare spectra in Fig. 2).

**Fig. 2.**
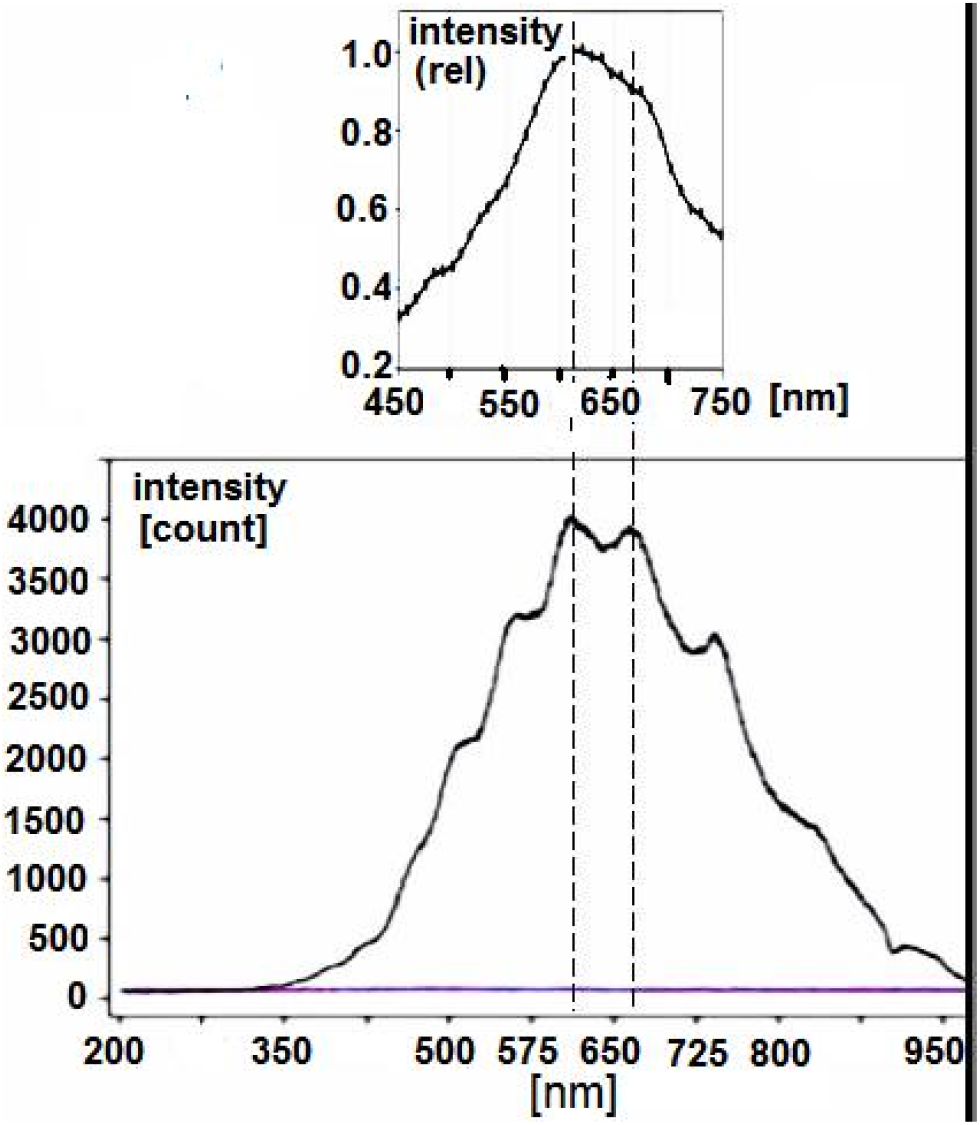
Correspondence of the spectrum of ultra-weak photon emission UPE from the ventral side of the human finger (top; according to M. Kobayashi et al. ^15^) and our protonic H^+^LED ^2^.

Considering protons as charge carriers, it can be concluded that, compared to electrons, the transport of protons in a biological (aqueous) environment is much more favorable than the flow of electrons (e.g. hydrated) ^22, 34^. The key is the ability to control the flow of protons owing to properties of proteins and protein complexes. Even single molecules such as amino acids are able to modify the aqueous environment in close proximity by creating an electrical potential barrier that is analogous to that in a typical electron-based p-n junction. Amino acid molecules are neutral, acidic or basic. They are able to modify water owing to interaction of proton-donor groups (acidic) and proton-acceptor groups (basic), transforming it into protonic analogues of n-type and p-type semiconductors, respectively ^6^.

The non-uniform electric field in the area of the formed protonic p-n junction is responsible for the controlled flow of protons ^4, 5, 6^. This enables protonic diodes and field effect transistors FETs to be formed in protein molecules ^4, 7, 8^. The same way, the complex protonic systems with an advanced functionality are created, e.g. the molecular motor C-ring of ATP synthase ^9-13, 23^ (Fig. 3b,c).

**Fig. 3.**
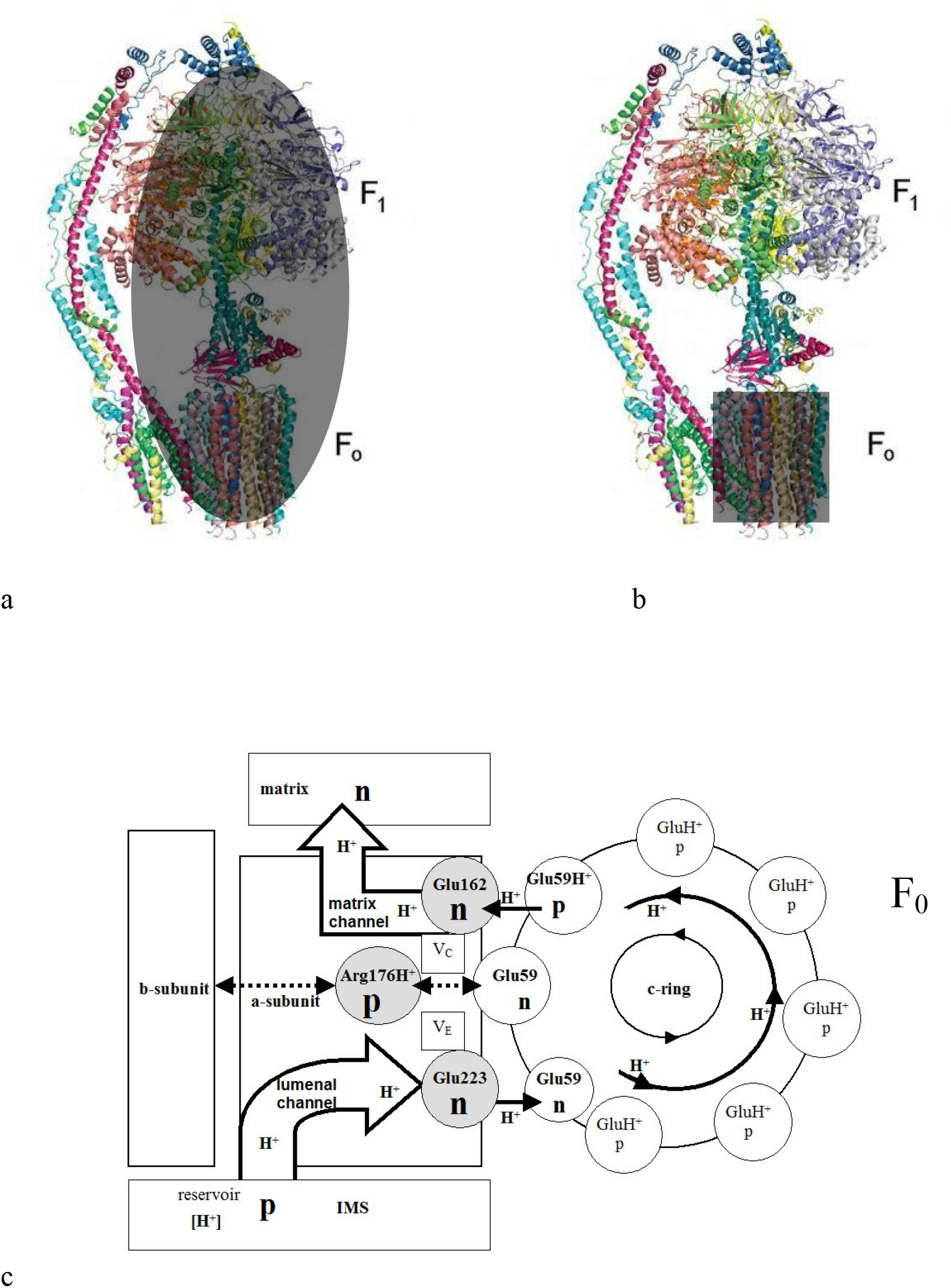
Marked grey F_0_F_1_ complex (a) and F_0_ unit (b) in ATP synthase/ATPase (a modified drawing based on publication ^18^). (c) The proton flow in the F_0_-ATP synthase (including C-ring) protonic p-n junction system ^37^ (a scheme based on the structure of ATP synthase/ATPase from ^18, 24, 27, 28, 29^). Preferred (“forward”) proton flow direction with possible light emission is from the “p” to “n” area, like in a protonic diode H^+^LED ^2, 4, 5^. We defined the protonic NPN transistor Glu162-Arg176-Glu223 ^4, 7, 8^, which is “normally off”, but even the small base current and low base-emitter voltage across Arg176-Glu223 will cause it to “turn on”, allowing large proton currents to flow collector (Glu223-Glu59-Glu162) and starting ATP synthesis. Protonic NPN transistors conduct when V_E_ is greater than V_C_. [This is discussed in a paper on the mechanisms of ATP synthase/ATPase powering and control based on the protonic p-n junction system (J. J. Langer at al., 2023; in preparation).]

We previously found that a protonic p-n junction in an aqueous medium is capable of generating light in the 350-1000 nm range with dominant emission in the visible (VIS) and near infrared (NIR) ranges, e.g. 550-750 nm with a maximum at 620 nm, when a proton current flows through it. Other tested H^+^LED devices emit mainly in the range of 620-800 nm with a maximum at 670 nm, but also in the range of 650-850 nm with a maximum at 750 nm ^2^.

This is particularly interesting in relation to biological systems in the context of biophotonics, as the proton current flowing through protonic p-n junction in proteins may induce emission of light ^2^. As expected, in this case the light is generated mainly at the molecular, sub-cellular level in structures with well defined proton flow, such as mitochondria ^23, 25^, leading to experimentally observed ultra-low photon emission (UPE) called bio photons. The emission is particularly intense (though still “ultra-weak” on a macroscopic scale) from structures with intense proton current, e.g. mitochondria, especially active ATP synthase and ATPase complexes, and in particular molecular motors, such as the C-ring of F_0_ unit ^14, 23, 25^.

The UPE spectrum discussed here covers the spectral range of 450-750 nm, with the region of dominant emission of the 570-670 nm and a main peak located in the 600-650 nm region, precisely with a maximum at 620 nm ^15^. The high adequacy between the H^+^LED and UPE spectra (Fig. 2) suggests that in both cases the emission mechanism is similar in terms of energetics and fundamental nature: the source of excitation is the flow of proton current through the protonic p-n junction. This is not surprising in light of the following.

### Modeling of ATPase emission

based on proton flow in the protonic p-n junction system of the C-ring F_0_ unit.

Considering the mechanism described previously, it can be expected that the flow of one proton through the protonic p-n junction can generate two excited water molecules and consequently - two photons (maximum) ^2^: H_3_O^+^ + -OH => 2 H_2_O* => h*ν* .

This means for ATP synthase (and ATPase), that at a speed of rotation of C-ring in F_0_F_1_ complex (Fig. 3a) of 300 s^-1^ and an average number of protons flowing per one rotation cycle of 1000 ^16, 17^, we have a cooperative proton current in C-ring of F_0_ unit ^29^ (Fig. 3b, 3c) of 3×10^5^ H^+^/s (4.8×10^-14^ A), which may generate total 6×10^5^ photons/s (maximum). Half of them (50%) can be observed at once due to the characteristics of the detection system (with a single detector light guide on one side), as used in experiments with H^+^LED ^2^). Thus, based on our model, we would expect to see the emission of about **3×10**^**5**^ **photons/s** (maximum) from a single ATP synthase/ATPase molecule when it is active.

### Simulation of ATP synthase/ATPase emission with H^+^LED

The protonic LED (H^+^LED), operated at 16 V and 0.8 A emits the light within the range of 350-950 nm with dominant emission in the range of 550-750 nm and main maxima at 620 nm and 670 nm ^2^. This is very similar to UPE within the range of 450-750 nm, with a dominant emission at 570-670 nm and the maximum at 620 nm (Fig. 2). In this case (H^+^LED), the current of 0.8 A corresponds to a proton flow of 5×10^18^ H^+^/s and the light intensity registered at 620 nm is of 4000 counts/s (164000 photons/s, calculated with an instrumental sensitivity ^20^ of 41 photons per count at 600 nm) at the aperture AP_1_ (the light emitting area, Fig. 4) of 0.19625 cm^2^ (∼0.2 cm^2^).

**Fig. 4.**
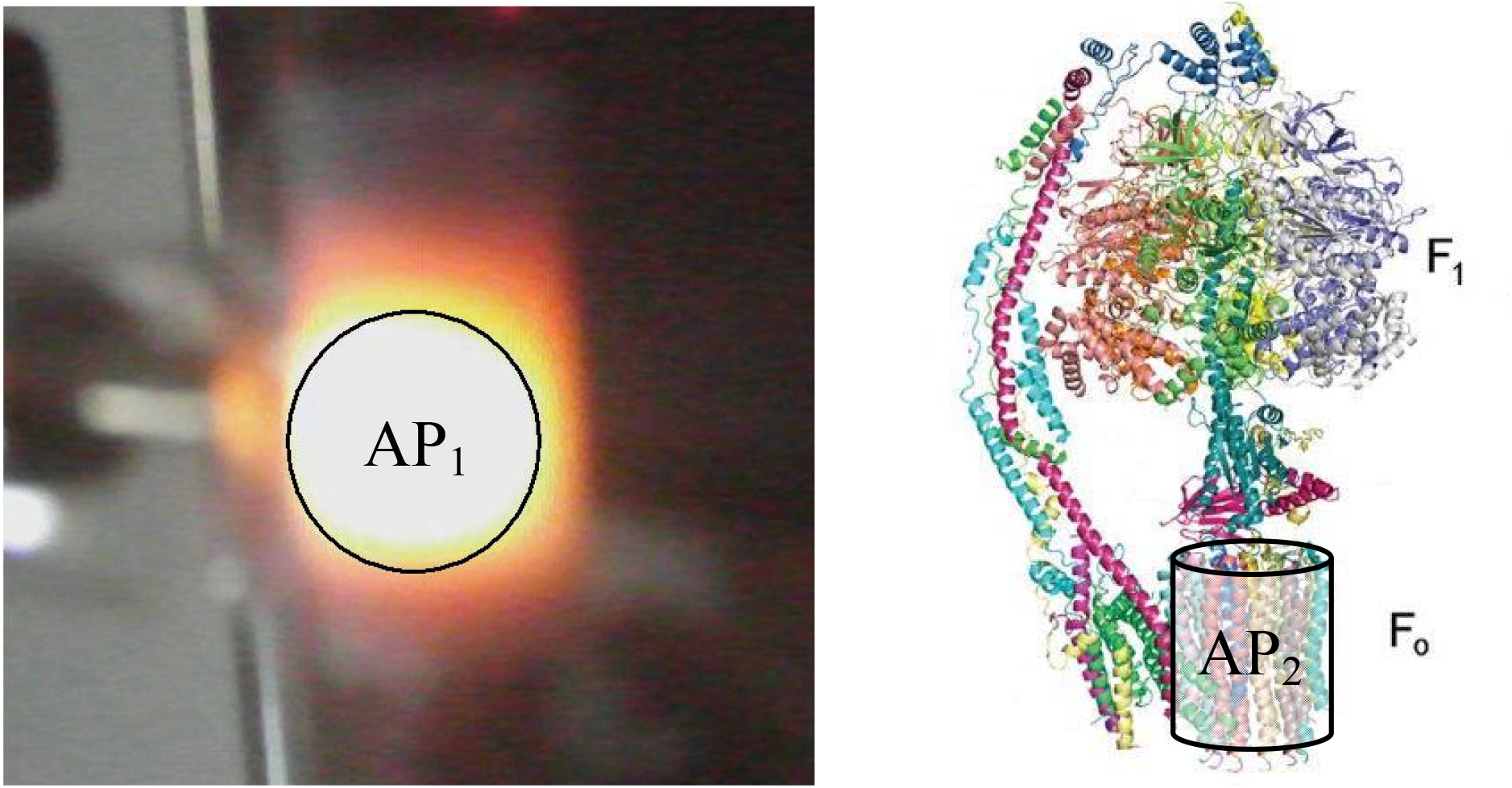
AP_1_ (H^+^LED) and AP_2_ (F_0_ ATP synthase/ATPase aperture definitions (a modified drawing based on publication ^18^).

With these parameters, the H^+^LED would emit 9.84×10^-9^ photons/s from the same light-emitting area (AP_1_ aperture, Fig. 4 left) with a proton flow of 3×10^5^ H^+^/s (4.8×10^-14^ A) calculated for the single C-ring F_0_ of ATP synthase/ATPase (the model in Fig. 3c).

However, the emission comes in fact from a single ATP synthase/ATPase molecule - F_0_F_1_ complex (Fig. 3a), and mainly F_0_ unit at ATP synthase of a maximum proton flow density (Fig. 3b, 3c) with an active (observable) emitting area of about 0.8×10^-14^ cm^2^ (aperture AP_2_, half of the side surface of the cylinder F_0_, Fig. 4).

Thus, to achieve the flux of photons of 9.84×10^-9^/s expected at an aperture AP_1_ of about 0.2 cm^2^, emitted from a source of an aperture AP_2_ corresponding in size to the F_0_ unit ^19^, the number of photons emitted from such source (of F_0_ unit) should amount to about <u>**2**.**4×10**^**5**^**/s**</u>.

This is proportional to the ratio of corresponding apertures AP_1_/AP_2_ equal 2.4×10^13^ and the macroscopic H^+^LED emission of 9.84×10^-9^ photons/s expected to be registered at the aperture AP_1_ for the proton flow of 3×10^5^ H^+^/s (from our model of ATP synthase/ATPase, Fig. 3c).

The result of simulation based on the H^+^LED experiment is unprecedentedly consistent with the ATP synthase emission theoretical modeling: of <u>**3×10**^**5**^ **photons/s**</u> (maximum), despite the difference in a basic structure of H^+^LED and ATP synthase/ATPase, because of a key role of protons and a protonic p-n junction in generating the light in both cases, just like described for H^+^LED ^2^. In addition, the simulation more accurately defines the molecular emitter, which corresponds in size to the C-ring of F_0_ ATP synthase unit (Fig. 3b, 4), the part with the highest proton current density in an active form. Thus, one can conclude that F_0_ units are the main source of bio photons (UPE) in living things, and UPE is powered by proton flow in F_0_’s protonic p-n junction system (like this presented in Fig. 3c) ^23^. At this point, the essential role of water in the functioning of the F_0_ motor unit should be emphasized. Water is necessary for formation the proton conduction channels ^23, 27, 28^, protonic p-n junctions ^4-7^ and the light emission (Fig. 5) ^2,3^.

**Fig. 5.**
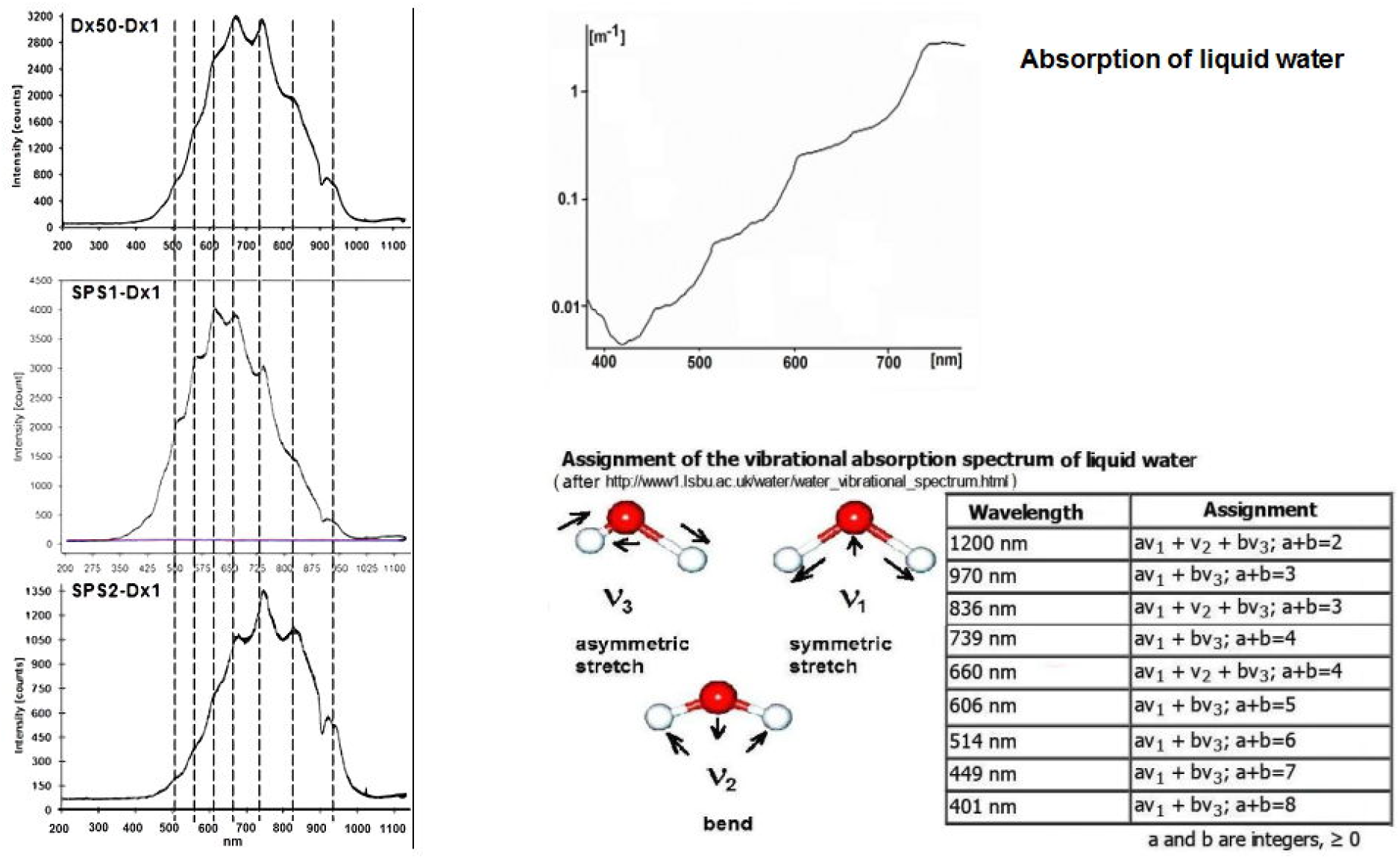
The emission spectra of protonic H^+^LEDs with different compositions (left): Dx50-Dx1, SPS1-Dx1 and SPS2-Dx1, show “fringes” associated with the same molecular vibrations in liquid water ^2^. Vibrational absorption of liquid water (right) ^34, 35^.

When the emission of bio photons at higher photon energies (above 3 eV, i.e. with a wavelength of 200-420 nm, corresponding to electronic excitations in water, Fig. 5) is mainly due to radicals and reactive oxygen species (ROS) ^1, 21, 32, 36, 37, 38^, bio photons with lower energy (below 3 eV, with a wavelength of 420-1000 nm) should be associated with the excitation of the protonic system (Fig. 5) as a result of the flow of the proton current (discussed in this work).

The correlation discussed here can be extended to other UPE spectra (Fig. 6) by considering the spectra of other H^+^LEDs, e.g. already published ^2, 3^, they all are related to molecular vibrations in liquid water and cover the spectral range of the UPE (350-1000 nm), Fig. 5. The emission is intense enough on the molecular and cellular scale (at a short distance from the emitter) to play an essential role in signaling and even local energy transfer, while being ultra weak on the macroscopic scale (UPE) due to light flux reduction and increase in light absorption with distance.

**Fig. 6.**
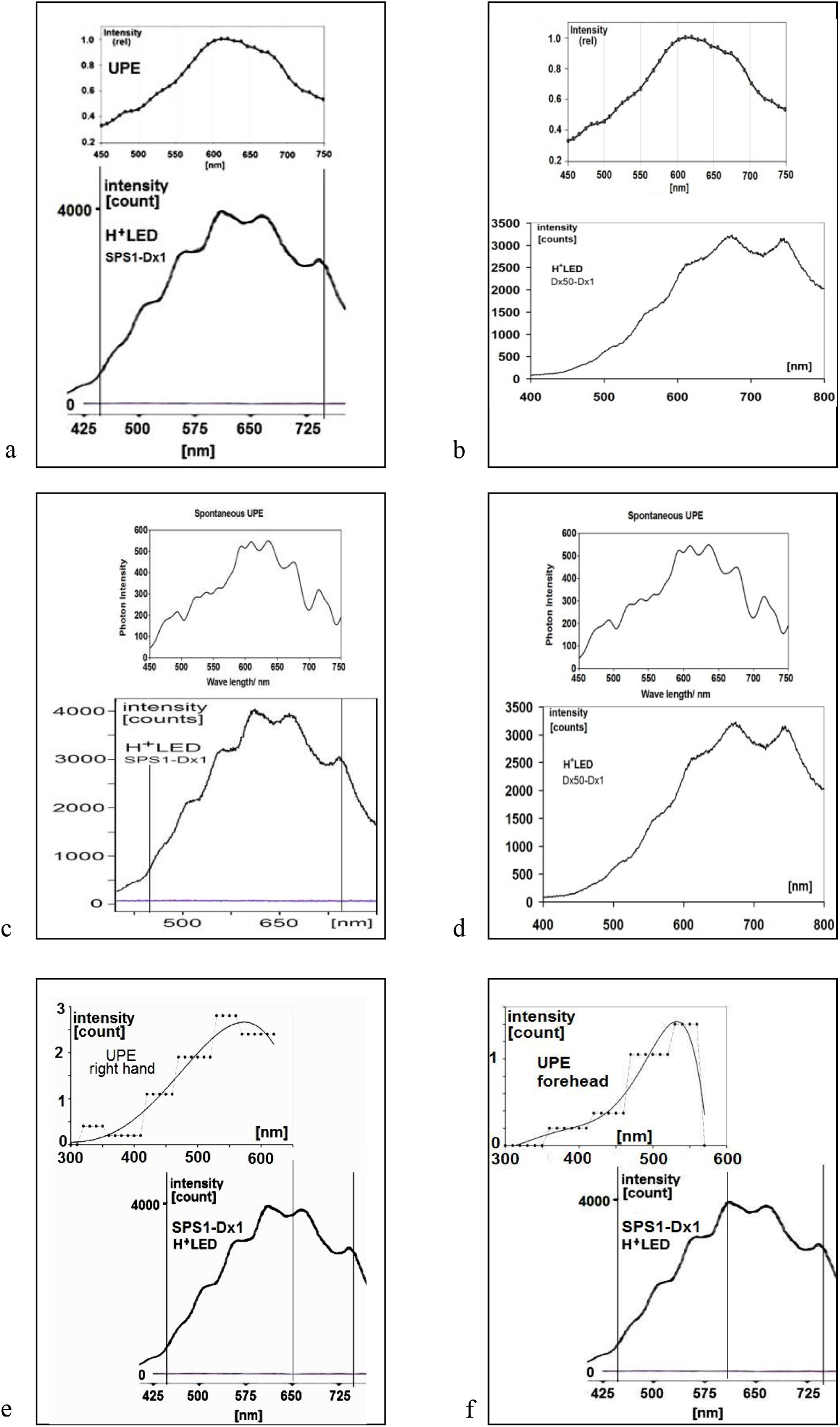
Examples of ultra-weak photon emission spectra UPE corresponding to protonic LEDs (H^+^LED ^2^) spectra: a) UPE from human body ^15^ and H^+^LED SPS1-Dx1 ^2^; b) UPE from human body ^15^ and H^+^LED Dx50-Dx1 ^2^; c) UPE from human body ^36^ and H^+^LED SPS1-Dx1 ^2^; d) UPE from human body ^36^ and H^+^LED Dx50-Dx1 ^2^; e), f) UPE from human body ^37^ and H^+^LED SPS1-Dx1 ^2^.

Taking this into account, one can expect that the UPE from individual anatomic locations would follow the pattern of mitochondrial density and dynamics within organs or tissues located nearby. Organs consuming greater amounts of energy produced by mitochondria more densely packed in their cells are more likely to emit greater amounts of bio photons. Thus, the activity of the brain containing hundreds to thousands of mitochondria in a single neuron ^30^, the heart containing more than 5,000 mitochondria in each muscle cell ^26^ or the liver containing more than 500 mitochondria per single hepatocyte ^31^, results in a greater intensity of UPE near the head, chest or stomach, as observed ^29, 32^. Responsiveness of touch receptor neurons has also been linked with increased mitochondrial density and shorter inter-mitochondrial distances ^33^, and this may explain significant contribution of UPE observed on hands ^32^. This supports our conclusions.

## Conclusions

Our finding explains why UPE (‘bio photons’) is only observed in living systems (organisms, tissues and cells) ^21^. This is due to the constant flow of protons in the active ATP synthase/ATPase ^23^ and in the mitochondria in general ^25-28^, which is necessary both for life and for the emission of light (bio photons).

In this context, a very specific flow of protons in the biological system of protonic p-n junctions (possible to be detected separately with NMR) generates bio photons (UPE), and both phenomena appear as the most fundamental biophysical markers of life at cellular and sub cellular levels.

In addition to scientific meaning, this results in important biomedical implications for diagnosis and therapy using UPE ^36, 37^.

The similarity of H^+^LED and UPE spectra (Fig. 2) allows the use of protonic H^+^LED as a new broadband light source, ideally suited to mitochondria-oriented low-intensity light therapy ^37^.

## Materials and methods

Water, an intrinsic protonic semiconductor, doped with an acid or a base forms protonic analogues of n-type or p-type semiconductors. Doping and the mechanical stability of the system are achieved using fictionalized polymers: ion exchangers Dowex 1×4 (**Dx1**) and Dowex 50W (**Dx50**) and sulfonated cross-linked polystyrenes, **SPS1** and **SPS2**, from our laboratory. The active groups are the acidic groups –SO_3_H from Dx50, SPS1 and SPS2, and the basic groups [N(CH_3_)_3_]^+^OH^-^ present in Dx1.

As a result of direct contact of wet materials with proton-donor and proton-acceptor properties, deposited in the form of layers between two Pt electrodes, a protonic p-n junction is formed, which exhibits rectifying properties and even light emission.

**Dx50** Dowex 50W(100–200 meshes) is a strongly acidic ion exchanger with functional groups - SO_3_H, used as is in the hydrogen form; matrix: styrene–divinylbenzene; cross-linked 8%.

**Dx1** Dowex 1×4 (100–200 meshes; SERVA) is a strongly basic ion exchanger, TYPE I, with functional groups (CH_3_)_3_–N^+^, used after activation with NH_4_OH; matrix: styrene–divinylbenzene (C_10_H_12_ . C_10_H_10_ . C_8_H_8_ . C_3_H_8_N)_x_, cross-linked 4%.

**SPS1** a strongly acidic sulfonated polystyrene, cross linked about 1.5%, [(C_8_H_8_-4C_8_H_7_SO_3_H).10H_2_O]_n_.

**SPS2** strongly acidic sulfonated polystyrene, cross-linked about 1% [C_8_H_7_SO_3_H.3H_2_O]_n_.

All information was described in detail in our previous paper [2].

## Author contributions

ML is responsible for collecting and analysing UPE data, including description and partial interpretation in the light of our concept; # research volunteering, cooperation with the Foundation for Friendly Education “Comet”.

JJL is the author of the overall concept, responsible for evaluating and interpreting the results and writing the manuscript;

## References

1. R. Van Wijk, E.P.A. Van Wijk, J. Pang, M. Yang, Y. Yan and J. Han, Integrating Ultra-Weak Photon Emission Analysis in Mitochondrial Research. Front. Physiol. 11:717, 1–15 (2020). doi: 10.3389/fphys.2020.00717

2. J. J. Langer, E. Frąckowiak, S. Golczak, Electrically induced light emission from proton-conducting materials. Protonic light-emitting diode. J. Mater. Chem. C, 8, 943—951 (2020). doi: 10.1039/C9TC05980F

3. J. J. Langer, E. Frąckowiak, Non-linear light emission of inorganic protonic diodes, H^+^LEDs, Advance Article, J. Mater. Chem. C, 9, 3052–3057 (2020). doi: 10.1039/D0TC05935H

4. J.J. Langer, Protonic p-n junction. Applied Physics A 34, 195 (1984).

5. J.J. Langer, A protonic rectifier diode. Applied Physics A 38, 59 (1985).

6. J.J. Langer, M. Martyń ski, Nano-scale protonic rectifier. Synth. Met. 107, 1–6 (1999).

7. J. Langer, Does protein form the molecular integrated electronic circuit based on protonic pn junctions? 14th International Symposium on the Chemistry of Natural Products (IUPAC), Poznań (Poland), 1984.

8. T. Miyake, M. Rolandi, Grotthuss mechanisms: from proton transport in proton wires to bioprotonic devices. J. Phys.: Condens. Matter 28, 023001 (2016).

9. P. Stevenson and A. Tokmakoff, Ultrafast Fluctuations of High Amplitude Electric Fields in Lipid Membranes. J. Am. Chem. Soc. 139 (13), 4743–4752 (2017). doi:10.1021/jacs.6b12412

10. D. Okuno, R. Iino, H. Noji; Rotation and structure of FoF1-ATP synthase, The Journal of Biochemistry,149(6), 655–664 (2011), 10.1093/jb/mvr049

11. M. Yoshida, E. Muneyuki & T. Hisabori, ATP synthase - a marvellous rotary engine of the cell. Nat Rev Mol Cell Biol 2, 669–677 (2001). 10.1038/35089509

12. J. Martin, J. Hudson, T. Hornung and W. D. Frasch, F_0_-driven Rotation in the ATP Synthase Direction against the Force of F1 ATPase in the FoF1 ATP Synthase. The Journal of Biological Chemistry, 290 (17), 10717–10728 (2015).

13. H. Guo, J. L Rubinstein, Cryo-EM of ATP synthases, Current Opinion in Structural Biology, 52, 71–79 (2018). 10.1016/j.sbi.2018.08.005.

14. M. Rahnama, J. A. Tuszynski, I. Bókkon, M. Cifra, P. Sardar, V. Salari. Emission of mitochondrial biophotons and their effect on electrical activity of membrane via microtubules. J Integr Neurosci. 10(1), 65–88, (2011). doi: 10.1142/S0219635211002622. PMID: 21425483.

15. M. Kobayashi, T. Iwasa, M. Tada, Polychromatic spectral pattern analysis of ultra-weak photon emissions from a human body. J. Photochem. Photobiol. B.159,186–90 (2016). doi: 10.1016/j.jphotobiol.2016.03.037. Epub 2016 Apr 7.

16. H. Ueno, T. Suzuki, K. Kinosita, Jr., and M. Yoshida, ATP-driven stepwise rotation of FoF1-ATP synthase. PNAS, 102 (5) 1333–1338 (2005).

17. D. F. Blair and S. A. Lloyd, Charged residues of the rotor protein FliG essential for torque generation in the flagellar motor of Escherichia coli. Journal of Theoretical Biology 266 (4), 733–744 (1997).

18. A. P. Srivastava1, M. Luo, W. Zhou, J. Symersky, D. Bai, M. G. Chambers, J. D. Faraldo-Gómez, M. Liao, and D. M. Mueller, High-resolution cryo-EM analysis of the yeast ATP synthase in a lipid membrane. Science 360, 6389 (2018); doi:10.1126/science.aas9699.

19. R. A. Capaldi, R. Aggeler, P. Turina, S. Wilkens, 1994. Coupling between catalytic sites and proton channel in FiFo-type ATPases. Trends Biochem. Sci. 19, 284–289.

20. Ocean Optics Inc., USB2000 Fiber Optic Spectrometer, Installation and Operation Manual, Document Number 170-00000-000-02-1005.

21. M. Benfatto, E. Pace, I. Davoli, M. Lucci, R. Francini, F. De Matteis, A. Scordo, A. Clozza, M. Grandi, C. Curceanu and P. Grigolini, Biophotons - new experimental data and analysis, arXiv/papers/2305/2305.09524, 05/16/2023. https://arxiv.org/ftp/arxiv/papers/2305/2305.09524

22. Ph. Colomban, Vibrational characterization of the various forms of (solvated or unsolvated) mobile proton in the solid state. Advantages, limitations and open questions. Solid State Ionics 393,116187 (2023). 10.1016/j.ssi.2023.116187

23. T.V. Zharova,, V.G. Grivennikova, V.B. Borisov, F1Fo ATP Synthase/ATPase: Contemporary View on Unidirectional Catalysis. Int. J. Mol. Sci. 24, 5417 (2023). 10.3390/ijms24065417

24. Ch. Bai and A. Warshel, Revisiting the protomotive vectorial motion of F_0_-ATPase, PNAS, 116(39), 19484–19489 (2019)

25. X. Wang, X. Zhang, Z. Huang, D. Wu, B. Liu, R. Zhang, R. Yin, T. Hou, C. Jian, J. Xu, Y. Zhao, Y. Wang, F. Gao, H. Cheng, Protons trigger mitochondrial flashes. Biophysical Journal 111, 386–394 (2016). 10.1016/j.bpj.2016.05.052

26. X. Wang, X. Zhang, D. Wu, Z. Huang, T. Hou, C. Jian, P. Yu, F. Lu, R. Zhang, T. Sun, J. Li, W. Qi, Y. Wang, F. Gao, H. Cheng (2017) Mitochondrial flashes regulate ATP homeostasis in the heart, eLife 6:e23908 (2017). 10.7554/eLife.23908

27. A. Marciniak, P. Chodnicki, K. A Hossain, J. Slabonska, and J. Czub, Determinants of directionality and efficiency of the ATP synthase F_0_ motor at atomic resolution. The Journal of Physical Chemistry Letters 13(1), 387–392 (2022). doi: 10.1021/acs.jpclett.1c03358

28. S. Kubo and S. Takada, Rotational mechanism of F_O_ motor in the F-type ATP synthase driven by the proton motive force. Front. Microbiol. 13:872565 (2022). doi: 10.3389/fmicb.2022.872565

29. N. Mitome, S. Kubo, S. Ohta, H. Takashima, Y. Shigefuji, T. Niina, S. Takada, Cooperation among c-subunits of FoF1-ATP synthase in rotation-coupled proton translocation. eLife 11:e69096 (2022). 10.7554/eLife.69096

30. M. Rango, N. Bresolin, Brain mitochondria, aging, and Parkinson’s disease. Genes (Basel) 9(5), 250 (2018). doi: 10.3390/genes9050250

31. D.D. Esposti, J. Hamelin, N. Bosselut, R. Saffroy, M. Sebagh, A. Pommier, C. Martel, A. Lemoine, Mitochondrial roles and cytoprotection in chronic liver injury. Biochem Res Int. (2012). doi: 10.1155/2012/387626

32. R. Van Wijk, M. Kobayashi, E.P.A. Van Wijk, Anatomic characterization of human ultra-weak photon emission with a moveable photomultiplier and CCD imaging. Journal of Photochemistry and Photobiology B: Biology 83, 69–76 (2006). doi:10.1016/j.jphotobiol.2005.12.005

33. A. Awasthi, S. Modi, S. Hegde, A. Chatterjee, S. Mondal, E. Romero, G.R. Sure, S.P. Koushika, Regulated distribution of mitochondria in touch receptor neurons of C. elegans influences touch response. bioRxiv, (2020). doi:10.1101/2020.07.26.221523

34. M. F. Chaplin, Structure and properties of water in its various states, Encyclopedia of Water: Science, Technology, and Society, Ed. P.A. Maurice, Wiley (2019). doi: 10.1002/9781119300762.wsts0002

35. M. F. Chaplin, Water Structure and Science, https://water.lsbu.ac.uk/water/water_structure_science.html

36. F. Zapata, V. Pastor-Ruiz, F. Ortega-Ojeda, G. Montalvo, A. Victoria Ruiz-Zolle, C. García-Ruiz, Human ultra-weak photon emission as non-invasive spectroscopic tool for diagnosis of internal states – A review. Journal of Photochemistry and Photobiology B: Biology, 216, 112141 (2021). 10.1016/j.jphotobiol.2021.112141

37. J. Tafur, E.P.A. Van Wijk, R. Van Wijk, and P. J. Mills, Biophoton detection and Low-Intensity Light Therapy: A potential clinical partnership, Photomedicine and Laser Surgery, 28 (1), 23–30 (2010) S Mary Ann Liebert, Inc. DOI: 10.1089/pho.2008.2373

38. M. Nerudová, K. Červinková, j. Hašek, and m. Cifra, “Optical spectral analysis of ultra-weak photon emission from tissue culture and yeast cells”, in Society of Photo-Optical Instrumentation Engineers (SPIE) Conference Series 2015, vol. 9450. doi:10.1117/12.2069897.

